# Genetic characterization of AmpC and extended-spectrum beta-lactamase (ESBL) phenotypes in *Escherichia coli* and *Salmonella* from Alberta poultry

**DOI:** 10.1101/2020.08.11.246645

**Authors:** Tam Tran, Sylvia Checkley, Niamh Caffrey, Rashed Cassis, Chunu Mainali, Sheryl Gow, Agnes Agunos, Karen Liljebjelke

## Abstract

Horizontal gene transfer is an important mechanism which facilitates bacterial populations in overcoming antimicrobial treatment. In this study, a total of 120 *Escherichia coli* and 62 *Salmonella enterica* subsp. *enterica* isolates were isolated from poultry farms in Alberta. Fourteen serovars were identified among *Salmonella* isolates. Thirty one percent of *E. coli* isolates were multiclass drug resistant (resistant to ≥ 3 drug classes), while only about 16% of *Salmonella* isolates were multiclass drug resistant. Among those, eight *E. coli* isolates had an AmpC-type phenotype, and one *Salmonella* isolate had an extended-spectrum beta-lactamase (ESBL)-type β-lactamase phenotype. We identified both AmpC-type (*bla*_CMY-2_) and ESBL-type (*bla*_TEM_) genes in both *E. coli* and *Salmonella* isolates. Plasmids from eight of nine *E. coli* and *Salmonella* isolates were transferred to recipient strain *E. coli* J53 through conjugation. Transferable plasmids in above total eight *E. coli* and *Salmonella* isolates were also transferred into a lab-made sodium azide-resistant *Salmonella* recipient through conjugation. The class 1 integrase gene, *int1*, was detected on plasmids from two *E. coli* isolates. Further investigation of class 1 integron cassette regions revealed the presence of an *aadA* gene encoding streptomycin 3”-adenylyltransferase, an *aadA1a/aadA2* gene encoding aminoglycoside 3”-O-adenyltransferase, and a putative adenylyltransferase gene. This study provides some insight into potential horizontal gene transfer events of antimicrobial resistance genes between *E. coli* and *Salmonella* in poultry production.

## 1 Introduction

For decades, antimicrobial resistance (AMR) has been a global issue of grave concern. Understanding potential mechanisms and driving forces for dissemination of genes encoding antimicrobial resistance between bacteria will help reduce the prevalence of resistant bacteria and thereby reduce risks to human and animal health. Acquisition of new resistance genes occurs frequently and naturally among bacterial communities from humans, animals and environments as outlined in the model known as ‘the epidemiology of AMR’ (John F. Prescott, 2006). However, the mechanism of dissemination of resistance genes is not yet fully understood.

*Escherichia coli* and *Salmonella* sp. are common bacterial causes of foodborne disease in human as well as gastrointestinal disease in animals (Folster et al., 2011; Ghodousi et al., 2015). *E. coli* is a genetically diverse species which has both commensal and pathogenic strains (Leimbach et al., 2013). *Salmonella enterica* are enteric pathogens, and are closely related to commensal *E. coli*, sharing ~85% of their genomes in common at the nucleotide level (Mcclelland and Wilson, 1998; McClelland et al., 2000).

AmpC-type CMY β-lactamase genes (*bla*_CMY_) have been found on both the chromosome and plasmids of many gram negative bacteria such as, *Klebsiella* sp., *Escherichia coli*, and *Salmonella* sp. CMY-2 is reported to be the most common plasmid-carried AmpC-type CMY in both *E. coli* and *Salmonella* isolates from various global regions including Asia, North America and Europe (Guo et al., 2014). Extended spectrum β-lactamases (ESBLs) are β-lactamases belonging mainly to Ambler class A, which includes TEM-, SHV-, CTX-M, GES, VEB enzyme families. ESBLs also include one enzyme family belonging to class D (Cantón et al., 2012). Isolates carrying plasmid-encoded AmpC can be easily misidentified as ESBLs due to their overlapping activity against beta-lactam antimicrobials. The inability to distinguish them could have significant treatment consequences (Hanson, 2003).

Mobile genetic elements, such as plasmids or DNA transposons, are the main mechanisms facilitating horizontal genetic transfer (HGT). Plasmid-mediated *bla*_CMY-2_ has been found to be the most predominant among other acquired *ampC* genes (Mata et al., 2012). The plasmids carrying *bla*_CTX-M_ or *bla*_CMY_ β-lactamase genes have been associated with transferable replicon types IncA/C or IncI1 (Hopkins et al., 2006; Guo et al., 2014).

Biofilm formation is the mode of growth of complex bacterial communities encased in exopolymeric matrix (O’Toole and Kolter, 1998). Biofilms not only increase bacterial tolerance to antimicrobial agents, but also create an intermediate environment for gene transferability (Molin and Tolker-Nielsen, 2003). Therefore, the impact of biofilm formation must be considered when assessing increased prevalence and persistence of AMR bacteria.

Antimicrobial use in the poultry industry improves animal health, welfare and production by preventing and treating animal disease resulting in lowered mortality, but may lead to the selection of AMR organisms (Diarra and Malouin, 2014). In Canada, the preventive use of ceftiofur in broiler chicken was banned in 2014 shortly after the surveillance of antimicrobial use and antimicrobial resistance in broiler chickens flocks in 2013 was implemented by the Canadian Integrated Program for Antimicrobial Resistance Surveillance (CIPARS) (Agunos et al., 2017). The most commonly used antimicrobial class in 2013 was coccidiostats, followed by bacitracin, virginiamycin and avilamycin (Agunos et al., 2017).

The isolates used in this study were a subset of those described in the 2015 CIPARS annual report (Public Health Agency of Canada, 2015). The aim of this study was to conduct further analysis of AMR phenotypes and characterize genetic mechanisms of AMR, not done previously.

## 2 Materials and Methods

### 2.1 Sampling, bacterial isolation and isolates used in this study

Fecal samples were taken from a single production unit on each of 30 registered premises/establishments (farms) participating in the CIPARS poultry surveillance in Alberta in 2015. Participating sentinel veterinarians were responsible for enrolling farms and collecting samples. Farms were chosen based on the veterinary practice profile and using specific inclusion and exclusion criteria. Samples were collected pre-harvest (broilers >30 days of age), approximately one week prior to slaughter. Fecal samples consisted of 10 fecal droppings from each of the four quadrants of the chosen barn/floor, pooled to represent the chosen production unit. This work was performed by Public Health Agency of Canada (PHAC) (Public Health Agency of Canada, 2015; FoodNet Canada, 2017).

A total number of 120 *E. coli* and 62 *Salmonella* isolates were isolated, banked, and shipped to the University of Calgary frozen on dry ice, by the Agri Food Laboratories Section of Alberta Agriculture and Forestry (Public Health Agency of Canada, 2015; FoodNet Canada, 2017). *E. coli* J53 (KACC 16628), a recipient isolate for the conjugation experiment, was received from the Korean Agricultural Culture Collection (KACC), Agricultural Microbiology Division, National Academy of Agricultural Science. *E. coli* HB101carrying plasmid pRK600, used as a helper strain, was received from the Dong lab, University of Calgary.

### 2.2 Susceptibility tests

Minimal Inhibitory Concentrations (MICs) of various antimicrobial agents were determined using Sensititre™ (TREK Diagnostic Systems, Inc.) antimicrobial susceptibility panels (CMV3AGNF) as part of the CIPARS (Public Health Agency of Canada, 2015). The same panel of antimicrobial agents was used for both *E. coli* and *Salmonella* isolates (Table 1 & Table 2). Antimicrobial resistance assays were conducted by the National Microbiology Laboratory (NML) St. Hyacinthine, and NML Guelph (Public Health Agency of Canada, 2015).

**Table 1.** AMR patterns in *Salmonella* isolates along with their serotypes. All the antimicrobials belonging to the same drug class were placed next to each other and separated from those in other drug classes by shading.

| *Salmonella*<br>serotype | Resistance pattern |  |  |  |  |  |  |  |  |  |  |  |  |  | Ratio |
| --- | --- | --- | --- | --- | --- | --- | --- | --- | --- | --- | --- | --- | --- | --- | --- |
|  | AMC | AMP | CRO | FOX | TIO | AZM | CHL | CIP | NAL | GEN | STR | SSS | SXT | TET |  |
| Enteritidis |  |  |  |  |  |  |  |  |  |  |  |  |  |  | 10/10 |
| Braenderup |  |  |  |  |  |  |  |  |  |  |  |  |  |  | 1/1 |
| Hadar |  |  |  |  |  |  |  |  |  |  |  |  |  |  |  |
| Pattern 1 |  | R |  |  |  |  |  |  |  |  | R |  |  | R | 6/10 |
| Pattern 2 |  | R |  |  |  |  |  |  |  |  |  |  |  | R | 1/10 |
| Pattern 3 |  |  |  |  |  |  |  |  |  |  | R |  |  | R | 3/10 |
| Hartford |  |  |  |  |  |  |  |  |  |  |  |  |  |  | 1/1 |
| Heidelberg |  | R | R |  |  |  |  |  |  |  | R |  |  | R | 1/1 |
| Infantis |  |  |  |  |  |  |  |  |  |  |  |  |  |  | 9/9 |
| Kentucky |  |  |  |  |  |  |  |  |  |  | R |  |  | R | 1/1 |
| Mbandaka |  |  |  |  |  |  |  |  |  |  |  |  |  |  |  |
| Pattern 1 |  |  |  |  |  |  |  |  |  |  | R | R |  | R | 1/4 |
| Pattern 2 |  |  |  |  |  |  |  |  |  |  |  |  |  | R | 3/4 |
| Schwarzengrund |  |  |  |  |  |  |  |  |  |  |  |  |  |  | 2/2 |
| Senftenberg |  |  |  |  |  |  |  |  |  |  |  |  |  |  | 5/5 |
| Thompson |  |  |  |  |  |  |  |  |  |  |  |  |  |  | 10/10 |
| Typhimurium |  |  |  |  |  |  |  |  |  |  |  |  |  |  | 4/4 |
| Worthington |  |  |  |  |  |  |  |  |  |  | R | R |  | R | 2/2 |
| I 6,7:k:- |  |  |  |  |  |  |  |  |  |  |  |  |  |  | 2/2 |
| Total |  |  |  |  |  |  |  |  |  |  |  |  |  |  | 62/62 |

There were seven drug classes tested: 1. β-lactam (AMC=Amoxycillin + Clavulanic acid, AMP= Ampicillin, FOX=Cefoxitin, CRO=Ceftriaxone, TIO= Ceftiofur), 2. Aminoglycosides (GEN=Gentamycin, STR=Streptomycin), 3. Quinolones (NAL=Nalidixic acid, CIP= Ciprofloxacin), 4. Sulfonamides (SSS= Sulfasoxazole, SXT= Trimethoprim sulfamethoxazole), 5. Macrolides (AZM= Azithromycin), 6. Phenicols (CHL=Chloramphenicol), 7. Tetracyclines (TET=Tetracycline). Drugs were grouped in classes by clear/grey areas.
R: Resistant

**Table 2.**
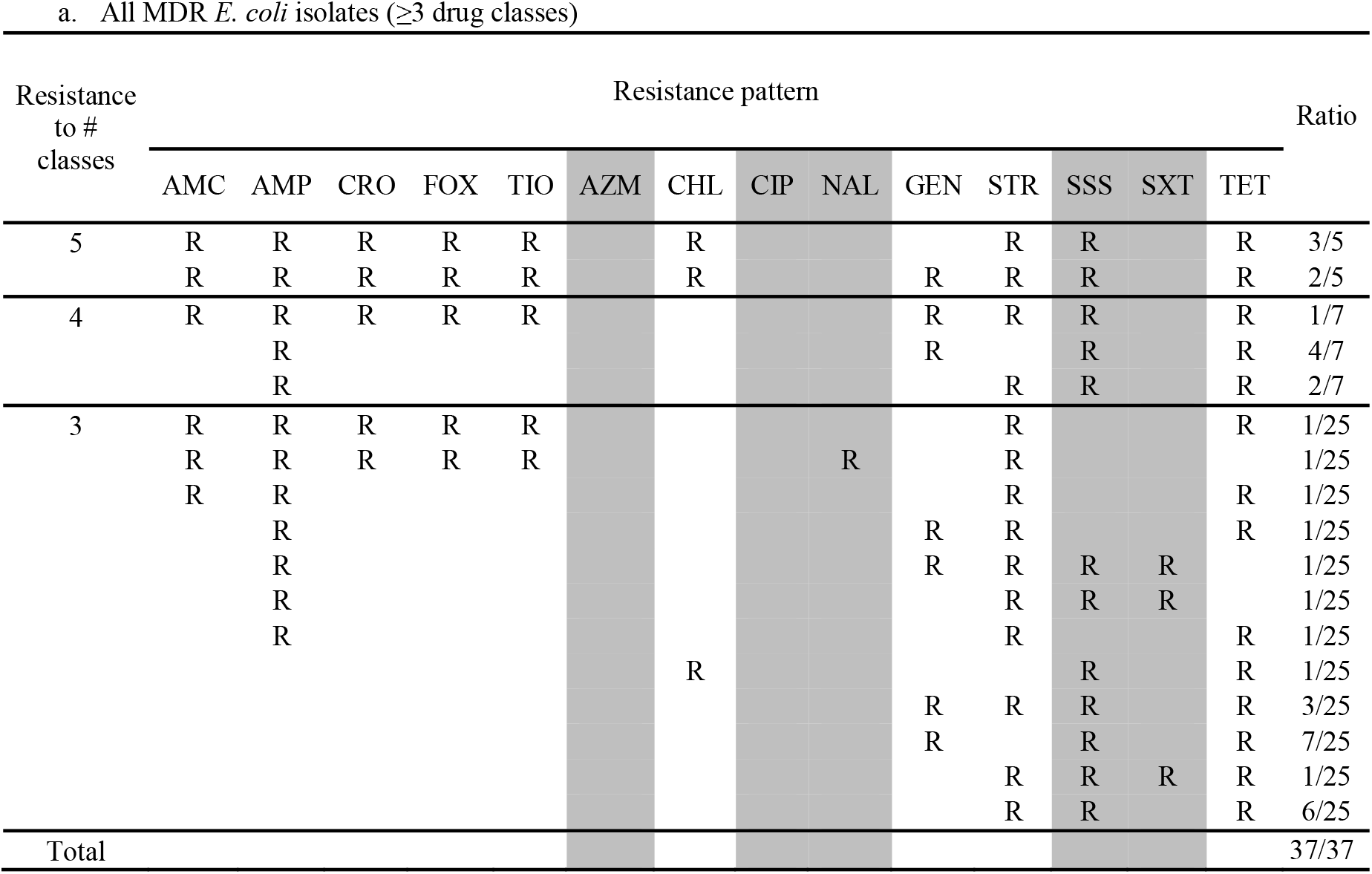
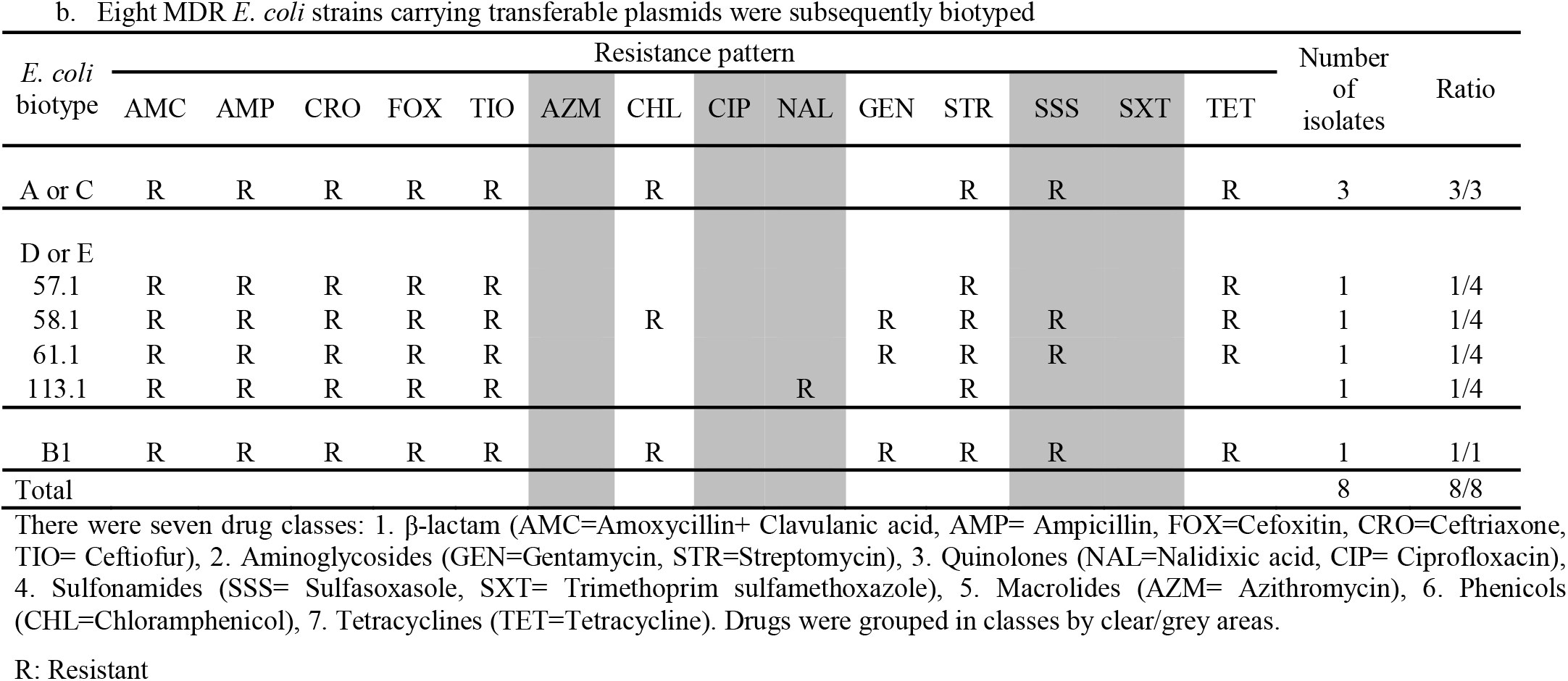
AMR patterns in MDR *E.coli* isolates. All the antimicrobials belonging to the same drug class were placed next to each other and separated from those in other drug classes by shaded or clear area.

The disc diffusion method was used to compare antimicrobial resistance profiles of isolate donors and *E. coli* recipients (Tendencia, 2004). Antibiotic discs were purchased from either BD BBL™ or Oxoid companies. The diameters of the zones of inhibition were recorded and interpreted according to Clinical and Laboratory Standards Institute (CLSI) guidelines (Clinical and Laboratory Standards Institute, 2013). For the purposes of this study, isolates displaying intermediate resistance were categorized as sensitive.

### 2.3 Phenotype and genotype confirmation of ESBL/AmpC genes

Two different ESBL/AmpC detection disc sets have been used to confirm ESBL/AmpC phenotypes. The first set is a combination of 4 individual discs of Cefotaxime/ Cefotaxime + Clavulanic acid/ Ceftazidime/ Ceftazidime + Clavulanic acid, purchased from either BD BBL™ or Oxoid company. The second set is a combination of 4 individual discs of Cefpodoxime/ Cefpodoxime + ESBL inhibitor/ Cefpodoxime + AmpC inhibitor/ Cefpodoxime + ESBL inhibitor + AmpC inhibitor, purchased from Mast Group company (D68C set).

In addition, AmpC and ESBL β-lactamase genes were detected using PCR assays. A total of three AmpC (*bla*_CMY-2_, *bla*_FOX_, *bla*_ACT-1/MIR-1_) and ten ESBL (*bla*_TEM_, *bla*_SHV_ *bla*_CTX-M-1_, *bla*_CTX-M-2_, *bla*_CTX-M-8_, *bla*_CTX-M-9_, *bla*_PER-1_, *bla*_VEB_, *bla*_IBC_/*bla*_GES_, *bla*_TLA_) β-lactamase genes were screened in AmpC/ESBL positive isolates. Primers used in the PCR assays are listed in Table 6.

The MDR *E. coli* and *Salmonella* were selected for further experiments after being confirmed to exhibit ESBL/AmpC phenotypes.

### 2.4 Plasmid characterization/ replicon typing

Plasmid miniprep was performed using an alkaline lysis method (Birnboim and Doly, 1984). Replicon typing was performed using PCR assay as described previously, with primers listed in Table 6 (Carattoli et al., 2005).

### 2.5 *E. coli biotypes/Salmonella* serovars

*E. coli* isolates of interest were assigned into one of four main phylogenetic groups by using a simplified two-step triplex polymerase reaction (Clermont et al., 2000). The results were confirmed using a quadruplex PCR assay which enabled us to classify isolates into a broader range of *E. coli* phylo-groups as well as distinguish them from the cryptic clades II to V (Clermont et al., 2013).

An assay to classify *Salmonella* serovars was performed by the PHAC serotyping laboratory as described previously (Public Health Agency of Canada, 2015).

### 2.6 Biofilm production assay *in vitro*

Isolates of interest were assessed for the ability to produce biofilm by culture in clear 96 well microtiter plates as described below (Stepanovic et al., 2007). An overnight liquid culture was diluted using a 1:100 ratio in tryptic soy broth (TSB) supplemented with casamino acids, and 200 μl of the suspension was then aliquoted onto a 96-well plate (each sample was assayed in triplicate). The plate was incubated as static culture at 37°C. After 24hr incubation, the culture medium was decanted and plate wells were rinsed with distilled water three times. The plate was stained with 200 μl of 1% crystal violet for 30 mins, rinsed under running tap water and then air-dried (by gently tapping then placing up-side-down on paper towels). Subsequently, 200 μl of glacial acetic acid (33%) was added into the wells to solubilize the dye, and absorbance was measured at 600 nm. The results were interpreted as described previously (Stepanovic et al., 2007).

### 2.7 Detection of integrons/integrases

To further study other mobile genetic elements, different integrase classes were identified using PCR assays (*int1, int2* and *int3*) using primers listed in Table 6 as described previously (White et al., 2001). Primers for amplifying the class 1 and class 2 integron cassette regions were used to detect the presence of resistance gene cassettes (Table 6) (White et al., 2001).

### 2.8 Conjugation experiment

Conjugation experiments were conducted using MDR isolates of interest as donors and *E. coli* J53 as the recipient with or without the presence of helper strain HB101/ pRK600. *E. coli* J53 (F-*met pro Azi*^1^), an *E. coli* K-12 derivative strain, is resistant to sodium azide (63). Recipient and donor strains were inoculated into LB broth and cultured overnight at 37°C. The next day, cells were harvested, washed with saline, and mixed together in a ratio of 1:1, and spotted on to LB plates. They were also spotted individually on LB plates as controls. After overnight incubation at 37°C, mating spots were washed and resuspended in saline; and different dilutions were plated on LB media containing sodium azide (0.2 gL^-1^) and ampicillin (100 μgml^-1^) to select transconjugants. Control spots were transferred to the same selective media to make sure that no growth was observed. Conjugation frequency was calculated by taking the ratio of the number of colonies counted on selective plates (LB supplemented with sodium azide (0.2 gL^-1^) + ampicillin (100 μgml^-1^)) for transconjugants over the number of colonies on selective plates (LB supplemented with sodium azide (0.2 gL^-1^)) for recipients. If there were no transconjugants obtained, a helper strain (HB101/pRK600) was added into the mating mix in the proportion of 1:1:0.5 (donor: recipient: helper strain) and spotted on LB plates as described. If there was no growth on plates selected for recipients, we added trypsin to the media to recover the recipients (42).

*Salmonella* isolate 112.2 were screened for spontaneously mutated colonies that resisted to sodium azide (Azi^R^) by plating on LB supplemented with sodium azide (0.2 gL-1). Then this Azi^R^ *Salmonella* was used as a recipient in conjugation with MDR isolates of interest as donors. Conjugation protocol was performed as described above.

### 2.9 Data visualization tools

Data visualization in this study was performed using following programs: Microsoft Excel 2013, R programming (R version 3.4.1).

## 3 Results

### 3.1 Sampling, isolation and identification of bacterial strains

Four *E. coli* isolates were obtained from each farm, resulting in 120 *E. coli* isolates from 30 farms. Twenty-three of 30 farms *Salmonella* positive, with between one and four isolates identified per farm. There were 14 different serovars identified among 120 *Salmonella* isolates (Table 1).

### 3.2 Antimicrobial susceptibility testing

Antimicrobial susceptibility in *E. coli* and *Salmonella* isolates is described. Isolates that were resistant to three or more drug classes were considered multiclass drug resistant (MDR). Thirty-one percentage of *E. coli* were MDR and 16% of *Salmonella* were MDR. About 4% of *E. coli* were resistant to five drug classes, while none of *Salmonella* were resistant to five drug classes.

The majority of *Salmonella* isolates were resistant to streptomycin and tetracycline (Table 1). There were 8 Salmonella serotypes that were sensitive to all tested drugs (Enteritidis, Typhimurium, Braenderup, Hartford, Infantis, Schwarzengrund, Senftenberg, Thompson). In addition to streptomycin and tetracycline, the majority of MDR *E. coli* showed resistance to sulfasoxasole (Table 2a). Among *E. coli* that were bio-typed, those belonging to groups D or E had diverse AMR patterns (Table 2b). Overall, *E. coli* isolates showed more diversity in resistance phenotype between farms than did *Salmonella* (Figure 1).

**Figure 1.**
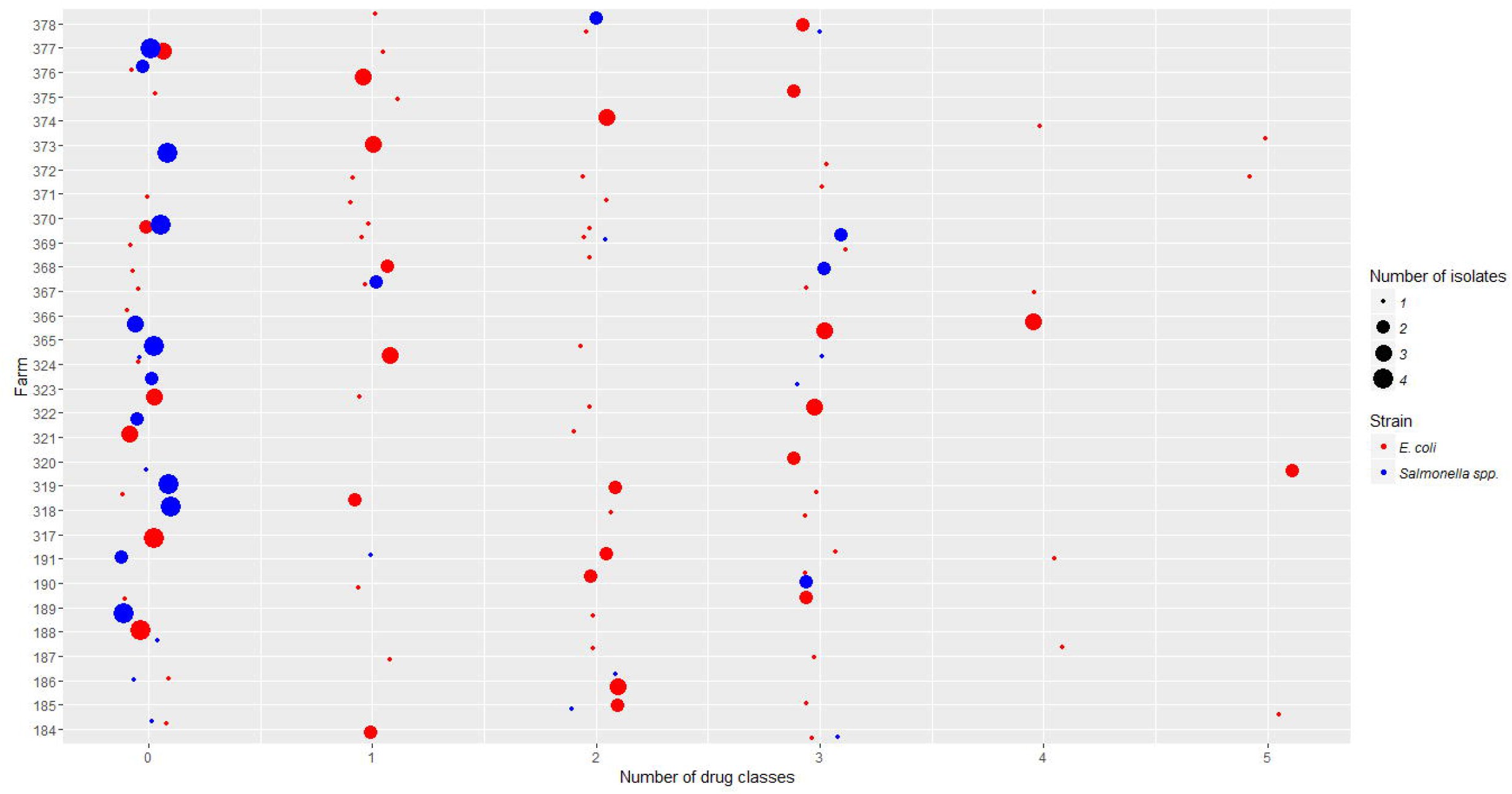
*Salmonella* isolates resistant to differing number of drug classes were compared to *E. coli* isolates across participating farms. The x-axis is the number of drug classes that isolates showed resistance to. The y-axis is the code of individual farms participating in this project. The size of each dot represents the number of isolates obtained in each farm. The color represents whether the isolates are *E. coli* or *Salmonella* strains.

Among MDR isolates, 19 *E. coli* and 10 *Salmonella* isolates with resistance to the β-lactam class of antimicrobials and at least two other drug classes, were selected for further study. These isolates came from 19 farms.

### 3.3 ESBL/AmpC phenotype and genotype

Eight out of 19 MDR *E. coli* isolates were resistant to both penicillin and cephalosporin β-lactam sub-classes and were confirmed as AmpC phenotype (Table 3). A unique *Salmonella* was resistant to both penicillin and cephalosporin sub-classes and was potentially an ESBL phenotype; in the presence of an ESBL inhibitor (clavulanic acid) the isolate showed sensitivity to penicillin (specifically amoxicillin). The ESBL phenotype was subsequently confirmed.

**Table 3.** Characteristics of MDR *E. coli* and *Salmonella* isolates showing ESBL/ AmpC phenotypes

| Strain | Isolate Number | Biotype<br>(*E.coli*)/Serotype<br>(*Salmonella*) | Plasmid<br>type | ESBL/ AmpC<br>phenotype | *bla*<sub>CMY-2</sub><br>gene | *bla*<sub>TEM</sub><br>gene | Integron/<br>Integrase | Biofilm<br>production<sup>c</sup> |
| --- | --- | --- | --- | --- | --- | --- | --- | --- |
| *E. coli* | 11.1 | A<sup>a</sup> / A or C<sup>b</sup> | A/C, I1 | AmpC | + | + | *aadA,aadA1a/A2* | + |
|  | 12.1 | A<sup>a</sup> / A or C<sup>b</sup> | A/C, I1 | AmpC | + | + |  | ++ |
|  | 57.1 | D<sup>a</sup>/ D or E<sup>b</sup> | A/C, I1 | AmpC | + | + |  | + |
|  | 58.1 | D<sup>a</sup>/ D or E<sup>b</sup> | A/C, I1 | AmpC | + | + |  | + |
|  | 61.1 | D<sup>a</sup>/ D or E<sup>b</sup> | A/C, I1 | AmpC | + | + | *int1, aadA* | + |
|  | 82.1 | B1<sup>a,b</sup> | A/C, I1 | AmpC | + | + | *int1* | + |
|  | 89.1 | A<sup>a</sup> / A or C<sup>b</sup> | A/C, I1 | AmpC | + | + |  | ++ |
|  | 113.1 | D<sup>a</sup>/ D or E<sup>b</sup> | A/C, I1 | AmpC | + | + |  | - |
| *Salmonella* | 119.2 | Heidelberg | A/C, I1 | ESBL | + | + |  | + |

<sup>a, b</sup> *E. coli* isolates were assigned into different phylogenetic group using both triplex and quadruplex PCR methods as described in previous studies, respectively (Clermont et al., 2000, 2013).
<sup>c</sup> The results were interpreted as no biofilm producer (-), weak biofilm producer (+), moderate biofilm producer (++).

The nine isolates which either showed the AmpC or ESBL phenotype were screened by PCR for a variety of AmpC and ESBL genes. The *bla*_CMY-2_ gene, an AmpC-type gene, and the *bla*_TEM_ gene, an ESBL-type gene, were identified on plasmids from all nine *E. coli* and *Salmonella* isolates. The sequence of the *bla*_TEM_ gene identified in this study shared 100% identity with the sequence of the *bla*_TEM_-116 gene found in *E. coli* strain MRC3 (accession no. KJ923009.1)

### 3.4 Plasmid characterization

Nine isolates were found to carry I1 and A/C-type replicon plasmids (Table 3). The plasmids varied in size from approximately 7kb to larger than 20kb. Only two *E. coli* isolates, 58.1 and 61.1, carried small plasmids (<10kb), while the rest carried larger ones (≥20 kb).

### 3.5 *E. coli biotypes/Salmonella* serovars of ESBL/AmpC-positive isolates

Both PCR methods confirmed that none of the *E. coli* isolates belonged to the group B2 (a group with high potential for pathogenicity) (Table 3). Biotypes of *E. coli* identified in this study were A, B_1_ and D (triplex PCR) or A, B_1_, C, D and E (quadruplex PCR). A *Salmonella* MDR isolate exhibiting the ESBL phenotype was identified as Heidelberg serovar.

### 3.6 Biofilm formation

In our biofilm assay, only one *E. coli* isolate lacked the ability to produce biofilm, while the other eight isolates produced biofilm (Table 3). Two *E. coli* isolates, 12.1 and 89.1, demonstrated moderate production, the highest amongst the isolates assayed.

### 3.7 Detection of integrons/integrases

Plasmids from two *E. coli* isolates, from 2 different farms, carried the class 1 integrase gene *int1* (Table 3). The class 1 integron cassette region was also detected in two isolates by PCR (Table 3). Sequencing these products identified the aminoglycoside resistance genes, *aadA* encoding streptomycin 3”-adenylyltransferase, and *aadA1a/aadA2* encoding aminoglycoside 3”-O-adenyltransferase. Class 1 integron cassettes were amplified from plasmids isolated from *E. coli* isolate 82.1. The results were confirmed by blasting the sequence against NCBI database. The sequences matched the sequence of a putative adenylyltransferase found on a plasmid isolated from the *Salmonella* Heidelberg strain N418 (Accession no. CP009409).

### 3.8 Transfer of resistance genes by conjugation

Plasmids were mobilized from all but one *E. coli* isolate (Table 4). No growth was observed on selective plates of either the transconjugants or recipients when attempting to conjugate *E. coli* isolate 61.1. However, the conjugation experiment of this isolate was successful when the media were supplemented with trypsin to reverse the effect of colicin produced by the donors. Additionally, the *E. coli* isolate 113.1 required the presence of the helper *E. coli* strain HB101 carrying helper plasmid pRK600 to enable movement of the plasmid to recipients.

**Table 4.**
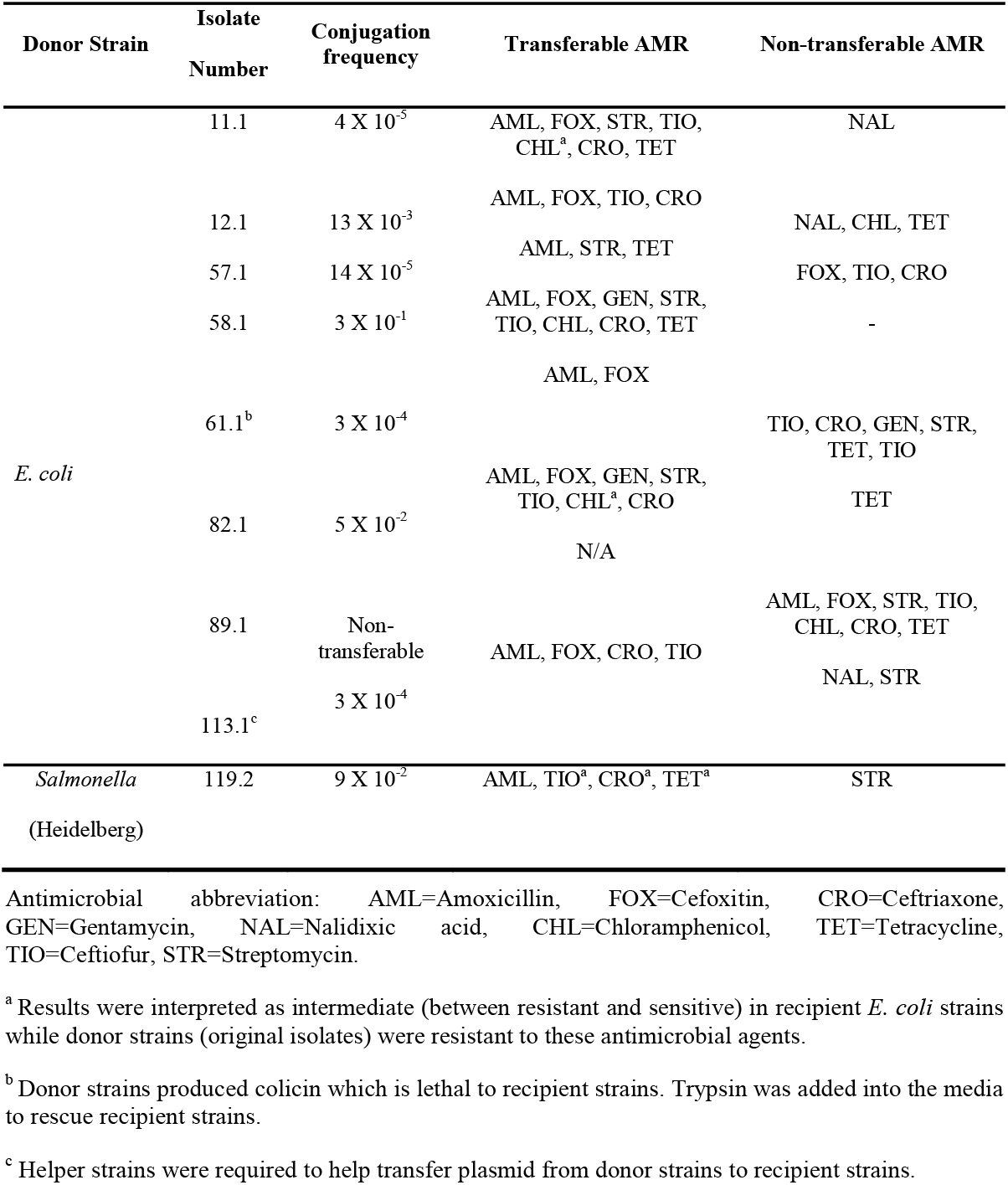
Conjugation frequency and AMR profile of transconjugants compared to donors (tested isolates) using *E. coli* J53 as a recipient

When using a lab-engineered sodium azide-resistant *Salmonella* as a recipient, we observed that plasmids from eight *E. coli* isolates with the exception of the isolate mentioned above, were able to move to *Salmonella* with variable conjugation frequency (Table 5).

**Table 5.** Conjugation frequency between donors (tested isolates) and a recipient strain (sodium azide-resistant *Salmonella*)

| Strain | Isolate ID<br>Number | Conjugation<br>frequency | Isolate ID<br>Number | Conjugation<br>frequency |
| --- | --- | --- | --- | --- |
| *E.coli* | 11.1 | $4 \times 10^{-9}$ | 61.1 | $5 \times 10^{-4}$ |
| | 12.1 | $1 \times 10^{-5}$ | 82.1 | $2 \times 10^{-2}$ |
| | 57.1 | $3 \times 10^{-7}$ | 89.1 | Non-transferable |
| | 58.1 | $7 \times 10^{-7}$ | 113.1 | $2 \times 10^{-4}$ |
| *Salmonella*<br>(Heidelberg) | 119.2 | $6 \times 10^{-2}$ | | |

**Table 6.**
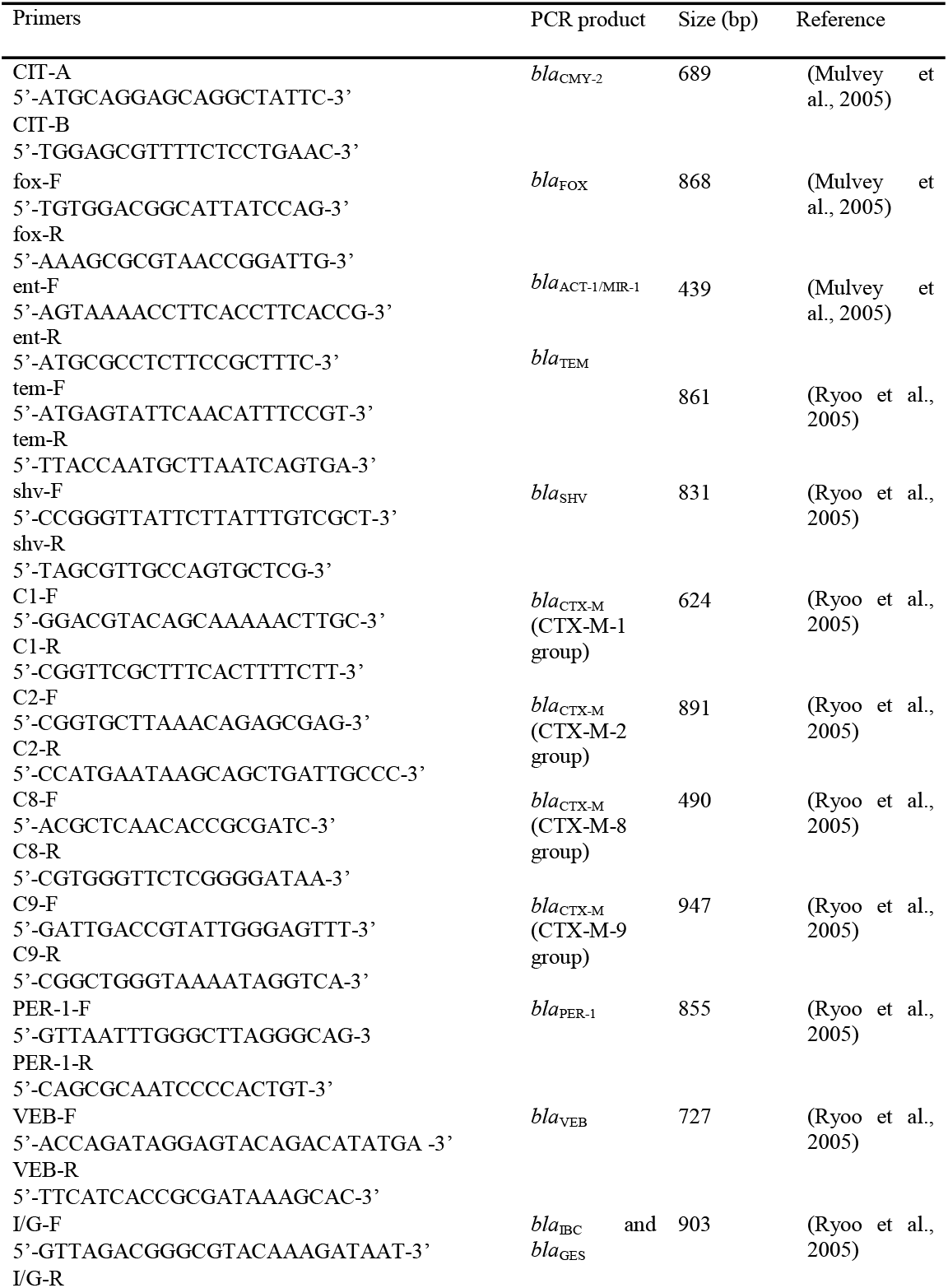
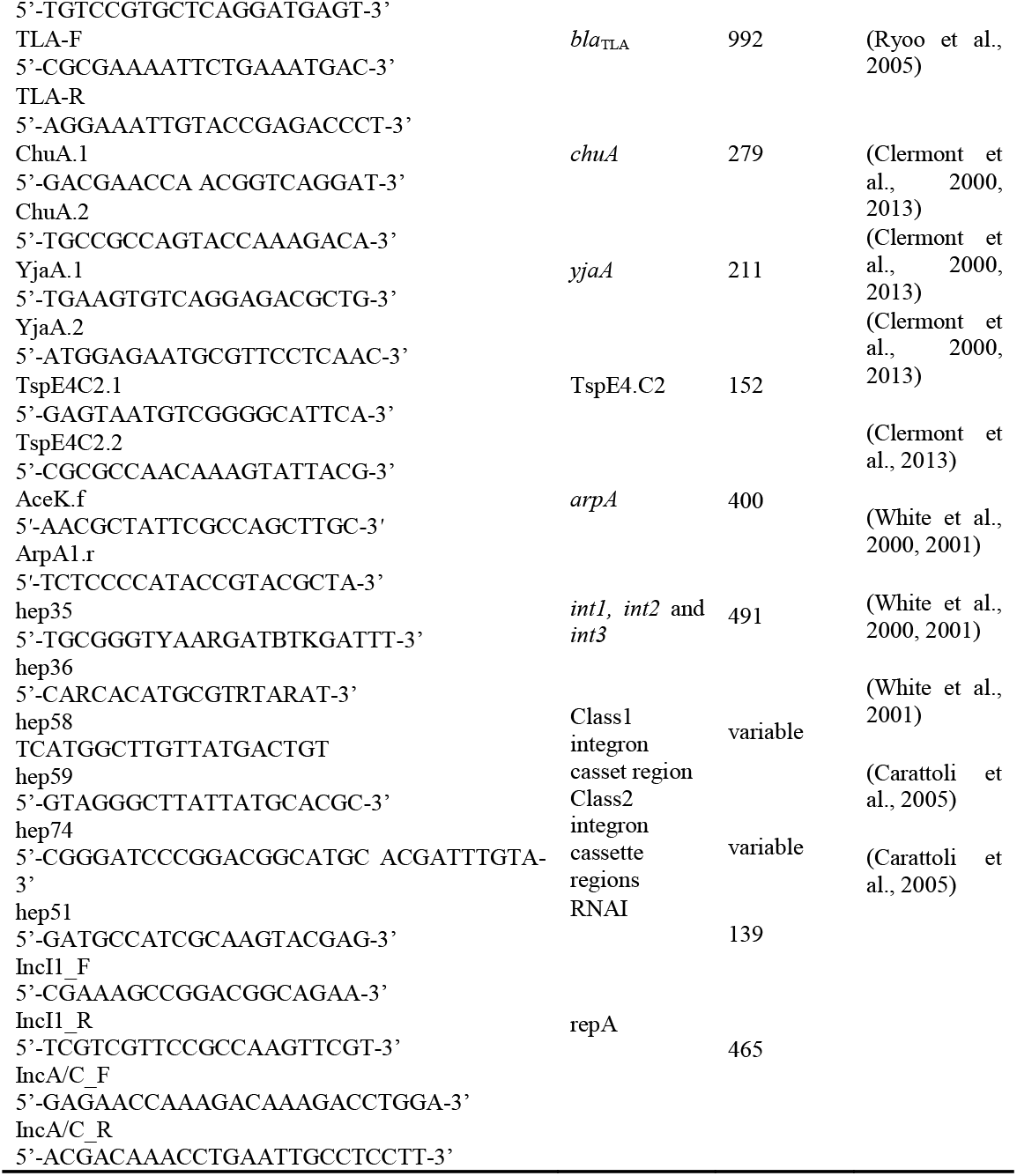
List of primers used in this study

### 3.9 Farm characteristics for nine isolates from which plasmids were mobilized

Seven ‘conventional’ (i.e., antimicrobials were used to some extent in all flocks) farms under the veterinary care of one practice were represented by the nine ESBL/AmpC phenotyped isolates in this study. All of the farms, excluding the farm providing isolate 89.1, received their chicks from the same hatchery. All birds were Ross 308 strain. The most frequently used antimicrobials were bacitracin and salinomycin administered in the poultry feed (n = 5), followed by the combination of penicillin and streptomycin administered in the water (n = 3). Avilamycin (n = 1), decoquinate (n = 1), monensin (n = 1), tylosin (n = 2) and the combination narasin and nicarbazin (n = 2) were also used as feed additives on these farms.

The number of chicks sampled per flock ranged from 14,790 to 55,000 within a single production unit. Age on the day of sampling ranged from 30 to 35 days old with an average weight ranging from 1.7 kg to 2.2 kg. The recorded floor space in the barns ranged from 8000 ft^2^ to 30550 ft^2^ and stocking density ranged from 0.54 to 0.67 ft^2^ per bird. Reported mortality rates ranged from 2.47% to 7.19% of the birds placed within the barn.

Hydrogen peroxide was used on three of the farms from which biofilm-producing bacteria were isolated, for cleaning of water lines between flocks. Five of the farms also used chlorine for treatment of their water lines during the production cycle. Footbaths (n = 3), dedicated farm clothes (n = 4) and gloves (n =2) were methods of farm biosecurity utilized. Manure was stored onsite in the vicinity of the barn on three farms. The most frequently reported method of cleaning the barns after each production cycle was washing only (n = 6) and chlorine products were used for disinfection on four farms.

## 4 Discussion

We examined the antimicrobial resistance phenotypes, genotypes, and mobile genetic elements of *E. coli* and *Salmonella* isolates isolated from chicken feces in Alberta farms in 2015. Thirty one percent of *E. coli* (37/ 120) were considered MDR, double the percentage of MDR *Salmonella* (16%, 10/62). Our study results are similar to those of a pilot study, which examined *E. coli* and *Salmonella* isolates in farming communities in northern Thailand, where *E. coli* had a higher diversity of antimicrobial resistance patterns compared to *Salmonella* (Hanson et al., 2002). Our study showed that a large portion of *Salmonella* isolates (71%) were susceptible to all 14 antimicrobials assayed. In contrast, only one-third of *E. coli* isolates (26%) were found to be susceptible to those 14 antimicrobials. About 4% MDR *E. coli* isolates were able to resist up to 5 drug classes.

The three most prevalent *Salmonella* serovars in our study were Enteritidis, Hadar, and Thompson. Serovars Typhimurium and Heidelberg were also identified. According to the National Enteric Surveillance Program (NESP) 2013 Annual Report, the three most commonly reported serovars in Canada, which has remained unchanged since 2008, were Enteritis, Heidelberg and Typhimurium (Government of Canada, 2015). Serovar Enteritidis is known as one of the most common *Salmonella* serovars found in poultry, and the second most prevalent cause of *Salmonella* infection in humans after the serovar Typhimurium (Suzuki, 1994; Porwollik et al., 2005; Trampel et al., 2014). Of 14 number of serovars in our study, only the Hadar serovar, was MDR in our study. The single only *Salmonella* serovar Heidelberg, was also the only with an ESBL phenotype, and the only MDR isolate. This is a of concern because in 2013-2014, a national outbreak of MDR *Salmonella* Heidelberg infections in the United States resulted in 200 hospitalized cases of 528 total cases (38%) (Gieraltowski et al., 2016). This outbreak was linked to chicken products from a single poultry company. In our study, six of the seven farms had chicks sourced from the same hatchery. There is potential for widespread dissemination of potentially virulent bacteria over a wide geographical region if such strains are present among eggs or chicks at the hatchery level. However, there was no evidence of this in our study

It was important to know if our MDR *E. coli* isolates that which exhibited either AmpC or ESBL phenotypes were also potentially pathogenic biotypes. PCR assays confirmed that none of these isolates belong to group B2, a group of virulent extraintestinal *E. coli*. Half of investigated isolates likely belong to group D, and are of lesser virulence. Phylogroups A and D were most frequently identified among our MDR *E. coli* isolates, similar to previous studies (Cortés et al., 2010; Coura et al., 2017).

Biofilm-producing bacteria tend to be more resistant to chemicals, detergents and antimicrobials. In addition, biofilm formation is known to be associated with up to 65 – 80% of all human clinical infections and is commonly implicated in chronic infections (Macia et al., 2014). Of the ESBL/AmpC-positive isolates assayed, eight of nine produced biofilm *in vitro*. Biofilm-forming ability might be a factor contributing to the dissemination of AMR genes among *E. coli* and *Salmonella* sp. isolates. Hydrogen peroxide and chlorine were used for treatment of water lines on some of the farms in question. It is possible that the use of these chemical methods alone, without the combination of some mechanical method of biofilm breakdown could lead to the selection and survival of MDR organisms in the poultry water lines.

It was important to know if our MDR *E. coli* isolates which exhibited either AmpC or ESBL phenotypes were also potentially pathogenic biotypes. PCR assays confirmed that none of these isolates belong to group B2, a group of virulent extraintestinal *E. coli*. Half of investigated isolates likely belong to group D, and are of lesser virulence. Phylogroups A and D were most frequently identified among our MDR *E. coli* isolates, similar to previous studies (Cortés et al., 2010; Coura et al., 2017).

Both *bla*_CMY-2_ and *bla*_TEM_ genes were present on plasmids isolated from in AmpC/ESBL positive MDR *E. coli* and *Salmonella* isolates. The same sequence of the genes, *bla*_CMY-2_ and *bla*_TEM_, was present in both *E. coli* and *Salmonella* isolates suggesting these genes might be circulating between them (Winokur, P. L., Vonstein, D. L., Hoffman, L. J., Uhlenhopp, E. K., & Doern, 2001). Interestingly, *E. coli* isolates showed the AmpC β-lactamase phenotype while *Salmonella* showed the ESBL β-lactamase phenotype. Even though they carried both *bla*_CMY-2_ and *bla*_TEM_ genes, only one of them was expressed in these *E. coli* and *Salmonella* isolates

The *bla*_CMY-2_ gene is the most common AmpC-type gene found in both *E. coli* and *Salmonella* from various sources: food, animals, and hospitals in multiple countries (Mulvey et al., 2005; Hiki et al., 2013; Cejas et al., 2014; Guo et al., 2014; Ghodousi et al., 2015). This gene has been hypothesized to have originated on the chromosome of *E. coli* and it could be induced with β-lactams in some Enterobacteriaceae such as *Enterobacter cloacae, Citrobacter freundii, Serratia marcescens*, and *Pseudomonas aeruginosa* (Sanders, 1987; Philippon et al., 2002). Unlike these bacteria, *E. coli* and *Salmonella* lack systems to produce inducible AmpC enzymes. Mutations in the *ampC* promoter have increased the resistance to oxyimino-cephalosporins in *E. coli* (Caroff et al., 1999).

Plasmids are considered to be facilitators for disseminating β-lactamase genes between various species such as *P. mirabilis, Achromobacter, Salmonella* and *E. coli* (Bobrowski et al., 1976; Levesque et al., 1982; Knothe et al., 1983; Bauernfeind et al., 1989). Molecular characterization of MDR plasmids is essential, yet complicated, because these plasmids are very diverse and promiscuous. The relatedness of plasmids can be analyzed using a PCR-based replicon typing method or whole genome sequence analysis (Carattoli et al., 2005). In previous studies, *bla*_CMY-2_-carrying plasmids found in either *E. coli* or *S. enterica* were most likely to belong to replicons I1 and A/C (Carattoli, 2009). Our results are also in accordance with these findings. *bla*_TEM_ genes have been reported to be located on plasmids of various replicon types such as A/C, I1, K, ColE, H12, etc (Carattoli, 2009). In our study, *bla*_TEM_ gene was found on plasmids of replicons A/C or I1 in one *Salmonella* isolate and eight *E. coli* isolates. Previously, multilocus sequence typing (MLST) showed plasmids encoding ESBL/AmpC genes in *E. coli* are highly promiscuous, resulting in the possibility of HGT between *E. coli* and related *Enterobacteriaceae* strains (Ewers et al., 2012).

In addition to plasmids, other mobile genetic elements including integrons, gene cassettes and integrase are also facilitating the spread of AMR genes (White et al., 2001). It was shown in multiple independent reports that there was a co-occurrence of integrase genes and AMR genes (Leverstein-van Hall et al., 2002; Marashi et al., 2012; Di Cesare et al., 2016). More specifically, a significant association was found between integrons and resistance to certain antimicrobials including gentamicin, kanamycin, streptomycin, tobramycin, sulfafurazole, trimethoprim, ampicillin, chloramphenicol, and tetracycline (White et al., 2001). In our study, the integrase gene *int1* was detected in two out of nine AmpC/ESBL-producing isolates. Using specific primers to amplify class 1 integron cassette regions revealed the presence of the aminoglycoside resistance genes *aadA* encoding streptomycin 3”-adenylyltransferase, *aadA1a/ aadA2* encoding aminoglycoside 3”-O-adenyltransferase, and a putative adenylyltransferase gene.

Conjugation *in vitro* showed that most of AmpC/ESBL positive MDR carried transferable plasmids that can disseminate AMR genes. Only the conjugation experiment on isolate 61.1 showed absolutely no growth on both transconjugant- and recipient-selective plates. We hypothesize that this isolate could have excreted some compounds that killed the recipients, such as colicin. To verify our hypothesis, trypsin was added into the media to recover the recipients. Previous studies have shown that treatment of cells with trypsin reversed the inhibition activity caused by colicin (Nomura and Nakamura, 1962; Dankert et al., 1980). Our findings confirmed that the isolate harbored a colicin-producing plasmid which prevented conjugal transfer of the R-plasmid unless the medium was supplemented with trypsin. This would likely prevent conjugal transfer in the natural microbial community as well. In addition, we were able to move plasmids between strains: *E. coli* ➔ *E. coli, E. coli* ➔ *Salmonella, Salmonella* ➔ *E. coli, Salmonella* ➔ *Salmonella*. By comparing conjugation frequencies, moving plasmids between *E. coli* and from *Salmonella* to *E. coli* recipients was much occurs at a higher frequency than moving plasmids from *E. coli* to *Salmonella*, or between *Salmonella*. We do not know if the same holds true in the natural environment.

In conclusion, we identified MDR isolates of *E. coli* and *Salmonella enterica* with ESBL/AmpC phenotypes, examined the sequences of the ESBL/AmpC genes in these isolates, and assayed their ability to form biofilm *in vitro*. We isolated and identified MDR plasmids which were readily transferred by conjugation between *E. coli* and *Salmonella* isolates. Results suggest the possibility of natural HGT by conjugation between *E. coli* and *Salmonella* may readily occur in the poultry house environment.

## Acknowledgements

We would like to thank the poultry veterinarians and producers who voluntarily participated in the CIPARS farm surveillance program and enabled data and sample collection. We are grateful to the Chicken Farmers of Canada and the Alberta Chicken Producers for their valuable input to the framework development and technical discussions.

This study was funded by Alberta Agriculture and Forestry (AAF) [grant number 2015R025R] with significant in a kind support from PHAC and the AAF, Agri-Food Laboratories.

## Author Contributions

TT conceptualized the research idea and experimental analysis. TT and NC designed and drafted the manuscript. NC did some statistical analysis. RG, CM, SG and AA did sampling and isolation work, performed antimicrobial testing and shipped isolates to our lab. SC wrote the proposal and applied for grants. KL and SC revised the drafted manuscript and made necessary corrections. The authors read and approved the final manuscript.

